# A multifaceted ecological assessment reveals the invasion of the freshwater red macroalga *Montagnia macrospora* (Batrachospermales, Rhodophyta) in Taiwan

**DOI:** 10.1101/2021.10.12.463998

**Authors:** Silvia Fontana, Lan-Wei Yeh, Shing Hei Zhan, Shao-Lun Liu

## Abstract

Invasive freshwater macroalgae are rarely described. Recently, a freshwater red alga, *Montagnia macrospora*, was introduced from South America to East Asia via the global aquarium trade. The earliest occurrence record of this alga in Taiwan is dated 2005. To determine whether *M. macrospora* has become an invasive species in Taiwan and to understand its traits that facilitated its invasion, we took a total-evidence approach that combines examination of ecological background and population genetic analysis. Our island-wide survey showed that *M. macrospora* alga was widespread in the field across Taiwan, where the climate greatly differs from that of South America. Our population genetic analysis revealed that the *cox2-3* sequences of all the specimens of *M. macrospora* from Taiwan were identical, consistent with the hypothesis that the alga expanded through asexual reproduction. Moreover, during our long-term ecological assessments and field surveys, we observed that *M. macrospora* is an ecological generalist that can self-sustain for a decade and bloom. Taken together, our data suggest that *M. macrospora* has successfully invaded the freshwater ecosystems in Taiwan due to its ability to disperse asexually and to grow under broad environmental conditions. We hope that our study brings attention to invasive freshwater algae, which have been overlooked in conservation planning and management.

## 1 INTRODUCTION

An alien species is a species introduced from its native range to a non-native range by human activities (e.g., Richardson et al., 2000). It can become naturalized (a self-sustaining founder population, but not spreading) in a non-native range and then may become invasive (overpopulated and/or widespread), potentially making adverse ecological impacts (e.g., wiping out native species and disrupting invaded ecosystems) and socioeconomic impacts (e.g., incurring conservation management costs) (reviewed in Richardson and Pyšek, 2012). Despite the enormous amount of effort and funding spent to limit the spread of invasive species, successful control and eradication of invasive species are extremely difficult (reviewed in Simberloff, 2021).

In freshwater environments, many introduced species of fish, vascular plants and invertebrates have been documented (reviewed in Havel et al., 2015). However, introduced species of freshwater macroalgae have received far less attention, with only a few known invasive cases — the green alga *Hydrodictyon reticulatum* in New Zealand (Hawes et al., 1991) and the red alga *Bangia atropurpurea* in North America (Kwei and John, 1977; Shea et al., 2014). The freshwater red alga *Montagnia macrospora* (Batrachospermales, Rhodophyta) is a recently reported introduced case. This alga is native to the tropical areas of South America, spanning French Guiana, Bolivia, and Brazil (Vis et al., 2008; Necchi et al., 2019). The earliest reported instances of *M. macrospora* in East Asia were found in a stream in Taoyuan in Taiwan in 2005 (Chou et al., 2014) and in an artificial pond in Okinawa, Japan in 2006 (Kato et al., 2009). In 2014, *M. macrospora* was abundantly found in 12 environments in Malaysia (Johnston et al., 2014). Recently, Zhan et al. (2021) found *M. macrospora* in many aquarium shops across East Asia, including Hat Ya (southern Thailand, Mayla Peninsula), Hong Kong, Taiwan, and Okinawa (Japan). A genetic analysis based on the plastid marker *rbcL* suggested that there were at least two independent introductions of *M. macrospora* into East Asia (Zhan et al., 2021). One of the introductions was revealed by a haplotype in aquarium shops in Taiwan and Japan, and the other introduction by a different haplotype in aquarium shops in Hong Kong and Thailand (Zhan et al., 2021). Thus, Zhan et al. (2021) proposed that *M. macrospora* was introduced into East Asia via the global aquarium trade. Although not traded as an ornamental organism, *M. macrospora* can hitchhike on ornamental organisms (e.g., aquatic plants), which are traded between South America and Taiwan (Zhan et al., 2021). It is unclear whether or not *M. macrospora* has become a naturalized species or an invasive species since its introduction into East Asia.

To assess the invasive potential of *M. macrospora*, it is crucial to understand its basic ecology. It is unknown whether *M. macrospora* is a generalist or specialist. Invasive species are typically generalists that are ecologically competent and can establish and spread their populations under a wide range of environmental conditions (e.g., light, temperatures, or nutrients) (reviewed in Gioria & Osborne, 2014). For example, introduced macroalgae can grow well in polluted waters with a high amount of certain nutrients (e.g., nitrate and phosphate), whereas native algae can typically grow well only in unpolluted waters (Inderjit et al., 2006).

Propagation via asexual spores or vegetative fragmentation is a fast and efficient way to spread in the absence of available mating partners. This ability can facilitate an alien species to self-sustain and thus to become naturalized in and invade non-native ranges. *M. macrospora* undergoes alternation between two free-living heteromorphic generations. As a haploid gametophyte (**Figure 1A**), the alga takes on a sexual thread-like mucilaginous form that produces and releases gametes (Necchi et al., 2019). As a turf-like diploid sporophyte (also known as chantransia; **Figure 1B**), the alga produces mostly asexual monospores and vegetative fragments (which can attach to other aquatic organisms and surfaces) and rarely sexual meiospores (Kato et al., 2009). In fact, the specimens of *M. macrospora* previously observed in a survey of Taiwan aquaria and streams were mostly spore-producing chantransia (Zhan et al., 2021).

**Figure 1.**
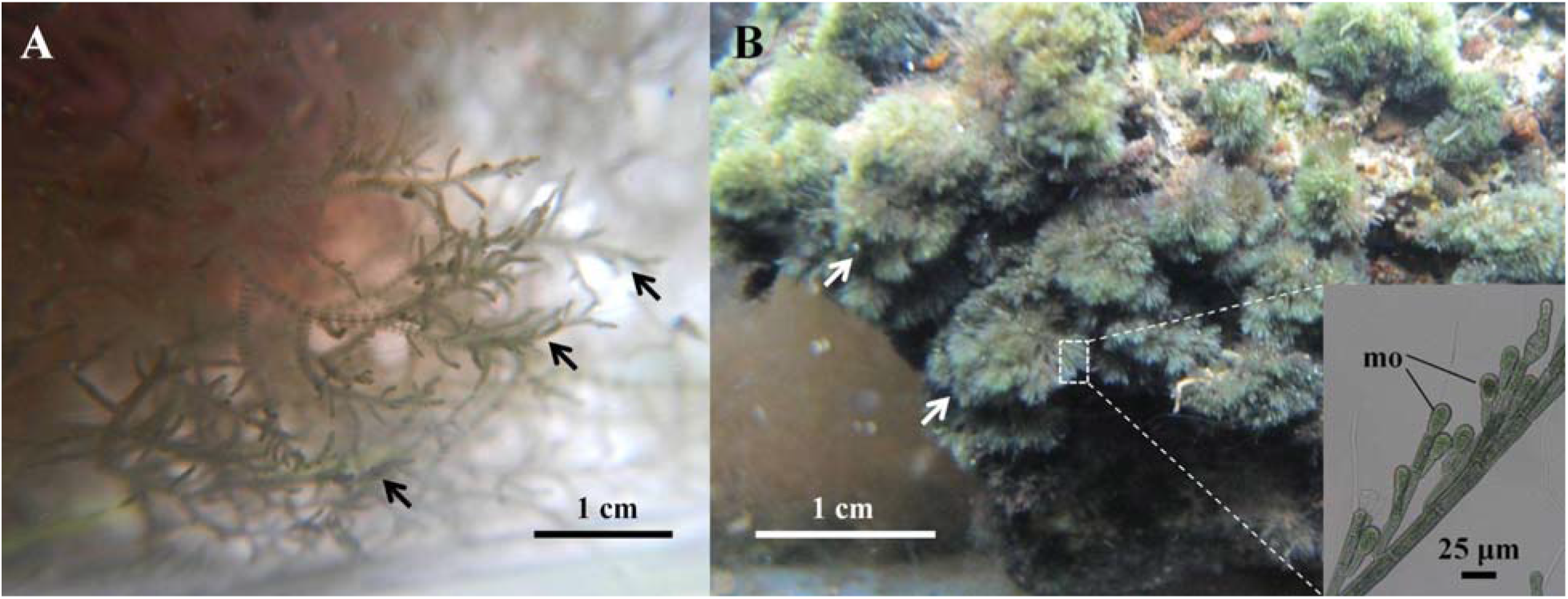
*In situ* photos of *Montagnia macrospora*. (A) A gametophyte (black arrows) collected from Maitaian, Hualien in eastern Taiwan. (B) Chantransia (i.e., sporophytes) (white arrows) from the Taoyuan stream, Taiwan. The inset indicates the close-up view of asexually reproductive structures - monospores (mo).

In this study, we took a total-evidence approach to determine whether *M. macrospora* has become an invasive species in Taiwan and to examine the traits of *M. macrospora* that might have facilitated its invasion. First, we inspected the presence of the alga to assess the extent of its spread across Taiwan. Second, we compared the niches of the alga in its native range in South America with the niches of the alga in its non-native range in Taiwan. Third, we determined the population structure of *M. macrospora* in South America and East Asia by reconstructing a haplotype network based on the intergenic spacer between the cytochrome oxidase subunit 2 and subunit 3 genes (*cox2-3*), which is widely used to evaluate inter-population genetic variation in freshwater red algae (e.g., Necchi and Vis, 2005, Paiano and Necchi, 2013, 2017). Fourth, we measured pH and the nutrient level of freshwater bodies around Taiwan where *M. macrospora* was found. Last, we performed a monthly ecological survey of a stream in Taoyuan, where Chou et al. (2014) reported the presence of *M. macrospora* in Taiwan for the first time in 2005. We collected data about the percent cover of *M. macrospora*, occurrences of other macrophytes, and water quality of the stream. This study provides a multifaceted assessment of the ecological status of *M. macrospora* as an invasive freshwater macroalgae in Taiwan. Filling this knowledge gap may be important to conservation planning and management for this invasive alga.

## 2 MATERIALS AND METHODS

### 2.1 Field inspections and assessments of pH and nutrient levels

To assess the distribution of *M. macrospora* in Taiwan, we carried out field inspections at 47 sites. *M. macrospora* was found in eight sites (**Figure 2**). We measured pH and the level of nutrients at seven sites. We were able to measure the level of nutrients at only seven sites, because we ran out of the necessary reagents. We also measured the level of nutrients in the artificial reservoir pond in Okinawa (Japan), since it was the place in East Asia where *M. macrospora* was first reported as an introduced freshwater macroalgae (Kato et al., 2009). The pH was measured using a pH meter (PH30, CLEAN Instruments Co., Taiwan). For nutrient measurements, water samples of 1,000 mL were collected and stored in an ice box during transportation. Then, measurements of four nutrients (ammonia, nitrate, nitrite, and phosphate, in mg L^-1^) were taken in the laboratory with a SMARTSpectro spectrophotometer (LaMotte, Maryland, USA).

**Figure 2.**
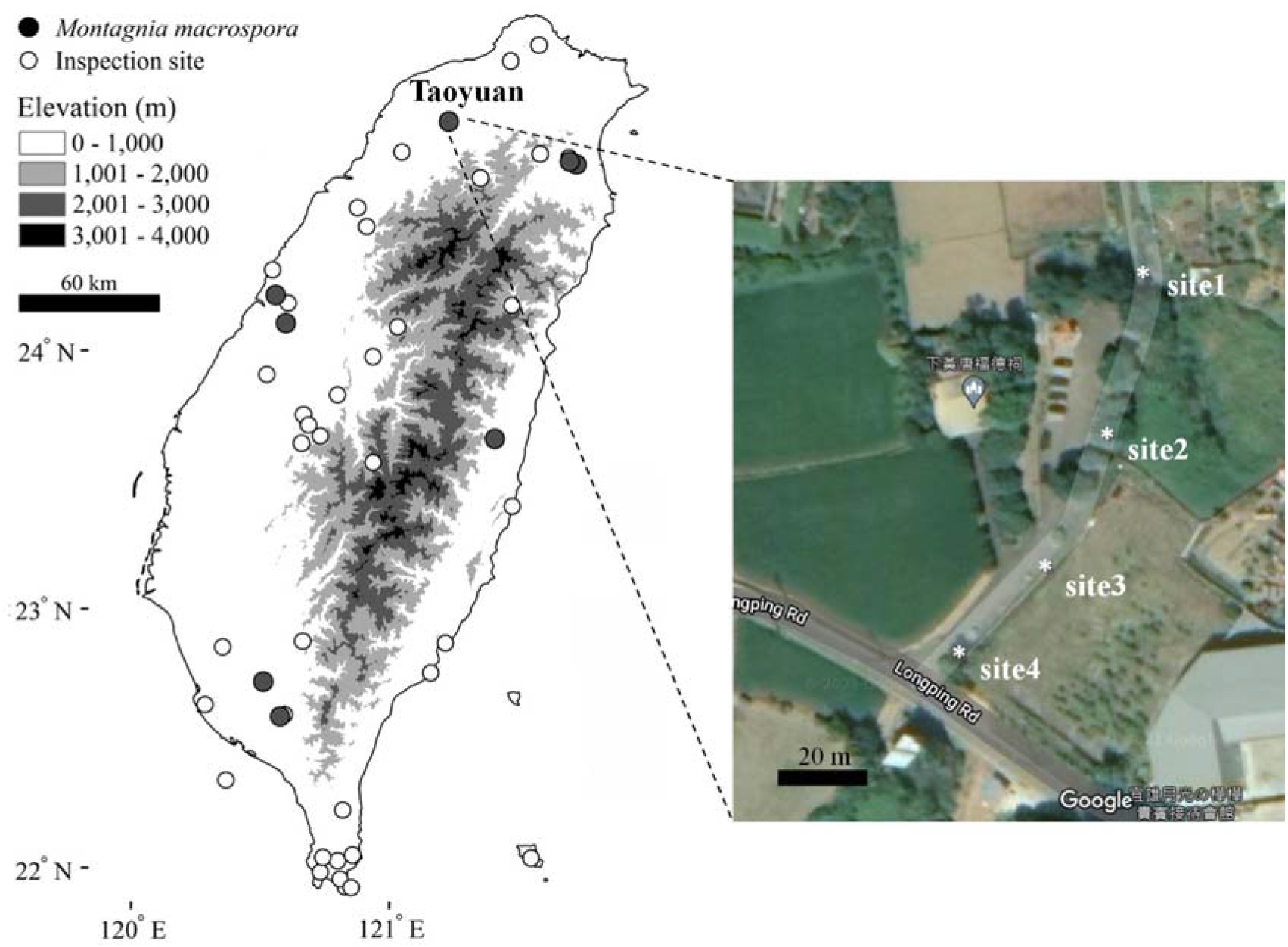
Sampling sites of *M. macrospora* in Taiwan. The grey circles mark the sites where *M. macrospora* was detected, while the white circles mark the sites where *M. macrospora* was not detected (as inspection sites). Google maps showing the monthly surveys at four sites, from site 1 (downstream) to site 4 (upstream), along the Taoyuan stream (24°53’09.3”N 121°13’54.1’’E).

### 2.2 Niche differences between the native ranges and the non-native ranges of *M. macrospora*

To assess the niche difference of *M. macrospora* between its native range in South America and its non-native ranges in Taiwan, we compared the climatic difference between these two areas. First, the current climatic CMIP5 data, which include 19 bioclimatic variables and elevation with a spatial resolution of 2.5 arc minutes, were obtained from the WorldClim database (http://www.worldclim.org/). To reduce cross correlation among the variables, we conducted a multicollinearity test using the variance inflation factor (VIF) using the R package usdm version 1.1-18 (Naimi et al., 2014). This resulted in the following ten predictors based on a VIF threshold of at least 10 (**Table S1**): elevation, mean diurnal range (bio02), isothermality (= mean diurnal range / temperature annual range * 100; bio03), mean temperature of wettest quarter (bio08), mean temperature of driest quarter (bio09), precipitation of wettest month (bio13), precipitation of driest month (bio14), precipitation seasonality (coefficient of variation; bio15), precipitation of warmest quarter (bio18), and precipitation of coldest quarter (bio19). The correlation between each variable never exceeded the recommended threshold of 0.75 (Kuman et al., 2006) (**Figure S1**). To identify the main climatic factors differentiating South America and Taiwan, we normalized the ten variables using the z-score approach and then applied a principal component analysis (PCA) using the R package stats version 3.6.0 (R Core Team, 2020). The sampling sites of *M. macrospora* were marked on a map of Taiwan using ArcGIS version 10.1 (ESRI, CA).

### 2.3 Phylogeographic analysis

Twenty-two specimens of *M. macrospora* were collected from aquaria and field sites (e.g., streams and springs) in Taiwan (**Table S2**). Two additional specimens were collected from Okinawa, where *M. macrospora* was introduced (Kato et al., 2009). A portion of each specimen (100 to 200 mg) was preserved in either silica gel or 95% ethanol for DNA analysis. Another portion of each specimen was suspended in a 10 to 15% formalin solution for morphological identification. Total genomic DNA was extracted using the ZR Plant/Seed DNA kit (Zymo Research, CA, USA), following the manufacturer’s recommendations. The genetic marker *cox2-3* was amplified using the primer pair: COX2F (5’-GTACCWTCTTTDRGRRKDAAATGTGATGC-3’) and COX3R (5’-GGATCTACWAGATGRAAWGGATGT C-3’) under the PCR conditions described in Vis et al. (2008). Sanger sequencing of PCR products was subsequently performed by the Mission Biotech Company (Taipei, Taiwan) as a service. The newly generated *cox2-3* sequence in this study was deposited in GenBank as the following accession number: OK442460.

For a phylogeographic analysis, we retrieved eight additional *cox2-3* sequences of *M. macrospora* specimens found in the native range of the alga in South America from Genbank as follows: EU106072-EU106079 (**Table S2**). The combined sequence set, which included our newly generated sequences and the sequences retrieved from GenBank, was aligned using MUSCLE (Edgar, 2004) via MEGA X version 10.1.7 (Kumar et al., 2018). Then, a haplotype network was inferred from the multiple sequence alignment using the TCS method (Clement et al., 2002), which takes a statistical parsimony approach, as implemented in PopART version 1.7 (Leigh and Bryant, 2015).

### 2.4 Ecological survey of a stream in Taoyuan, Taiwan

Field surveys over the 14 months between August 2012 and October 2013 were carried out monthly at four sites along a small spring-fed stream in Longtang District, Taoyuan City, Taiwan (24°53’09.3”N 121°13’54.1”E). The four sites were 10 m apart from each other, going from downstream (site 1) to upstream (site 4) (**Figure 2**). The stream ran adjacent to a small alley in a flat farm area. The bed of the stream was made of mud and silt, artificial substrate of cement and gravel, and some sandbags (**Figure S2**).

Each site showed variation in vegetation and surrounding structures, which caused differences in illumination. Site 3 enjoyed the most sunlight, and the other three sites showed different degrees of tree canopy shading. Site 1 was muddier than the other sites, sites 2 and 3 had cemented stream beds, and site 4 had silts in its bed (**Figure S2**). To calculate the percent cover of *M. macrospora*, five 10 x10 cm quadrats were equally spaced along the 80 cm-wide stream beds at each site. For each site, the percent cover of the algae was calculated as the average of the percent cover of the five quadrats of the site. *M. macrospora* was identified *in situ* based on the color (bluegreen) and overall external morphology of the thallus. This was followed by *cox2-3* molecular verification of five randomly collected specimens per site (data not shown). Light (μmol photons m^-2^ s^-1^), pH, conductivity (μS cm^-1^), turbidity (mg L^-1^), water velocity (m s^-1^), and water depth (cm) were measured *in situ* by using a portable digital photometer (DL-204 EZDO, Chi-Jui Instrument Enterprise, Taiwan), a pH meter (PH30, CLEAN Instruments Co., Taiwan), a meter for both conductivity and turbidity (CON30, CLEAN Instruments Co., Taiwan), a flow probe (FP111 Global Water, Smart Scientific Corp., Taiwan), and a ruler, respectively. For nutrient measurements, the level of four nutrients (i.e., ammonia, nitrate, nitrite, and phosphate) were determined as described above. Precipitation data at Zhongli automatic weather station, located in Zhongli District, Taoyuan City was retrieved from Central Weather Bureau, Taiwan (https://www.cwb.gov.tw).

We examined the variation in and relationships between the percent cover of *M. macrospora* and environmental variables. First, to determine whether there was any variation in the percent cover of *M. macrospora* and environmental variables among the four survey sites, we applied a one-way Analysis of Variance (ANOVA), followed by multiple comparisons among each pair using a *post hoc* Tukey test. Prior to ANOVA, the light intensity among the four sites of each month was normalized based on the relative light percent difference (i.e., corresponding to different degrees of light shading by tree canopy). Second, to examine the effects of the environmental variables on the percent cover of *M. macrospora* at each site, we applied stepwise multiple regression analyses using the R package MASS version 7.3-54 (Venables and Ripley, 2002). For this part of the statistical analysis, we did not include light, velocity, and water depth, because they were affected by rain clouds and rainfall during or before our field work. All the statistical analyses were performed in R (R Core Team, 2020).

In addition to the monthly survey of the stream in Taoyuan, we performed sporadic observations in the Taichung Industrial Park Outlet, and reported the percent cover of *M. macrospora*, pH and nutrients level from January 2014 to September 2021.

### 2.5 Co-occurring, non-native aquarium-associated macrophytes

During each survey in the Taoyuan stream, we recorded the presence of other aquarium-associated freshwater red algae and introduced aquatic plants that are commonly found in aquarium shops. Species identification of aquatic plants was performed based on several guidebooks (Lin, 2009a, b, c). Species identification of freshwater red algae was done based on *rbc*L sequences.

## 3 RESULTS

### 3.1 Occurrences across Taiwan and niche differences between Taiwan and South America

Our island-wide inspections revealed the presence of *M. macrospora* in eight out of 47 (~17%) sites in Taiwan (**Figure 2**). A PCA based on ten climatic variables indicated that the eight sites in Taiwan were different from the 16 sites in South America (**Figure 3**). The first two components (PCs) of the PCA accounted for 74.3% of the total variance of the data (PC1: 43.4% and PC2: 30.9%; **Figure 3**). The sites in Taiwan and the sites in South America were primarily differentiated by PC2 (**Figure 3**). We took the load threshold of the absolute value of 0.3 as a rule of thumb to determine which variable substantially contributed to PC2. The variables bio02 and bio03 were negatively loaded, whereas the variables bio08, bio13, bio15, and bio18 were positively loaded (**Table S3**). Compared with South America, the niches of the alga in Taiwan had lower mean diurnal range (bio02) and isothermality (bio03), but had higher mean temperature of wettest quarter (bio08), precipitation of wettest month (bio13), precipitation seasonality (bio15), and precipitation of warmest quarter (bio18). In other words, the niches of the alga in Taiwan are different from the niches in South America.

**Figure 3.**
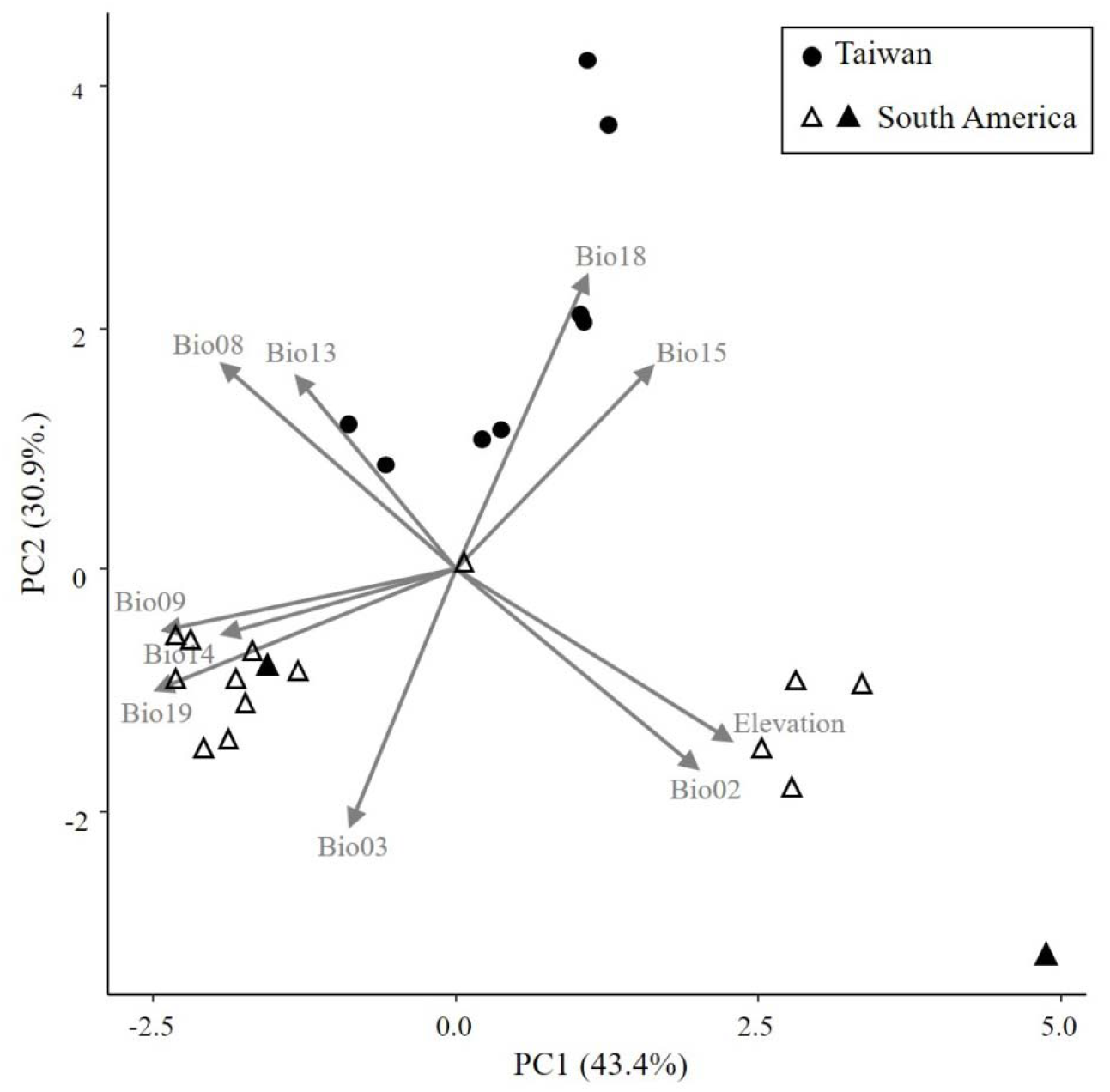
A principal component analysis showing the climatic differences between the non-native range of the alga in Taiwan and the native range of the alga in South America. Arrows indicate the degree of correlation between ten variables and two principal components (PC1 and PC2). The black circles mark the sites in Taiwan, while the white and black triangles mark the sites in South America. The black triangles represent the populations from Bolivia and French Guiana that have an *rbc*L haplotype identical to those in Taiwan.

### 3.2 Lack of nucleotide diversity in *cox2-3* in *M. macrospora*

The *cox2-3* sequence alignment from 32 specimens of *M. macrospora* was 338 bp in length. There was a 4 bp gap in all the sequences from Taiwan and Japan and in five out of the eight haplotypes from South America. The haplotype network analysis indicated identical *cox2-3* sequences in all the specimens collected from both aquaria and fields in Taiwan and Japan (**Figure 4**). In contrast, higher genetic diversity was found in the specimens of the algae from the native range in South America, consistent with the pattern of the *rbcL* haplotype diversity shown in Zhan et al. (2021).

**Figure 4.**
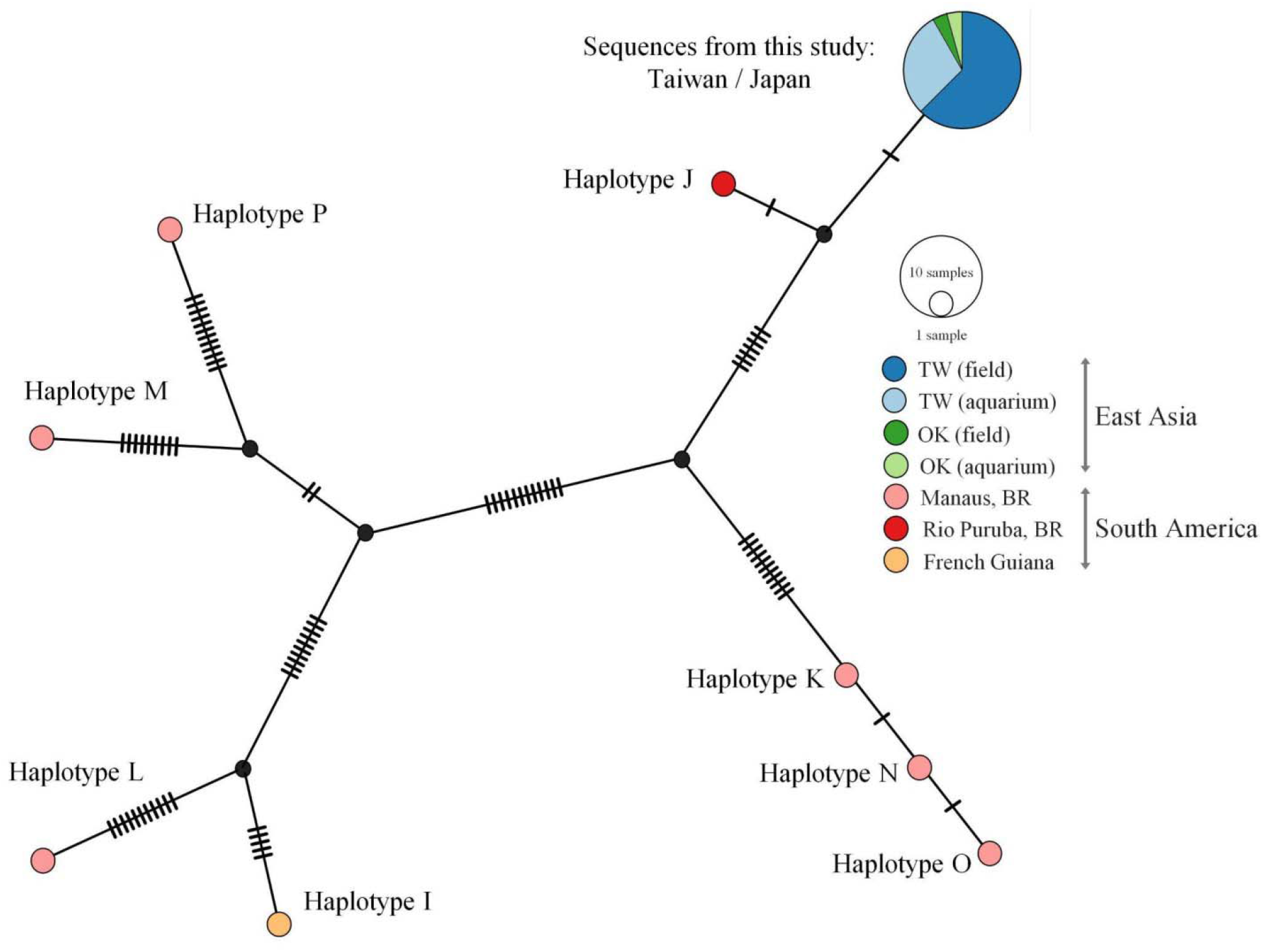
A haplotype network of *M. macrospora* inferred based on *cox2-3* using the TCS method. The hatch marks on the links between the haplotypes represent nucleotide differences between the two haplotypes. One haplotype represents the sequences obtained from this study (from Taiwan and Japan) and the other eight haplotypes belong to *M. macrospora* from South America (the sequences were retrieved from NCBI Genbank, keeping their names as in the original reference). The samples from South America show higher genetic diversity. None of the sequences from South America share the same haplotypes with the samples obtained from this study. Haplotype J, however, is the closest, with only two mutational steps of separation from the haplotype found in Taiwan and Japan.

### 3.3 Presence of *M. macrospora* in Taiwan in a wide range of pH and nutrient levels

We examined pH and the level of nutrients at the seven field sites in Taiwan and at one site in Okinawa where we observed *M. macrospora*. A broad range of pH was observed from the most acidic one (5.67) to the most basic one (8.34). In the Taoyuan stream, nitrate (> 5 mg L^-1^) was detected at a higher level than phosphate, ammonia, and nitrite (on average, 0.05, 0.31, and 0.09 mg L^-1^, respectively) (**Figure 5**). In other Taiwan locations, with the exception of Taichung Industrial Park Outlet, the levels of nitrate were lower than the Taoyuan stream, at 2.08 mg L^-1^ on average. We consistently detected higher proportions of nitrate compared to phosphate, ammonia, and nitrite (on average, 0.11, 0.5, and 0.15 mg L^-1^, respectively). A notable outlier among the samples measured was Taichung Industrial Park Outlet. At this location, the level of ammonia (146 mg L^-1^) and the level of phosphate (109 mg L^-1^) were ~300 times higher than the levels at the other locations in Taiwan. The levels of nitrate and nitrite at the Taichung Industrial Park Outlet (12 and 1.4 mg L^-1^, respectively) were also notably higher than the other locations in Taiwan (2 and 15 times higher than the level of the Taoyuan stream, respectively) (**Figure 5**). Kato et al. (2008) reported that the artificial reservoir pond in the Senbaru-ike in Okinawa, Japan, where *M. macrospora* was reported “very eutrophic” but did not take nutrient measurements to quantify how eutrophic the pond was. We visited the same pond in 2013 to measure the level of nutrients in the pond and confirmed that the pond was indeed more eutrophic than most of the sites in Taiwan (ammonia, 3.34 mg L^-1^; nitrate, 1.1 mg L^-1^; nitrite, 0.22 mg L^-1^; and phosphate, 0.79 mg L^-1^) (**Figure 5**). Our results showed that *M. macrospora* can grow under a wide range of pH and nutrient conditions, and tolerate highly eutrophic waters.

**Figure 5.**
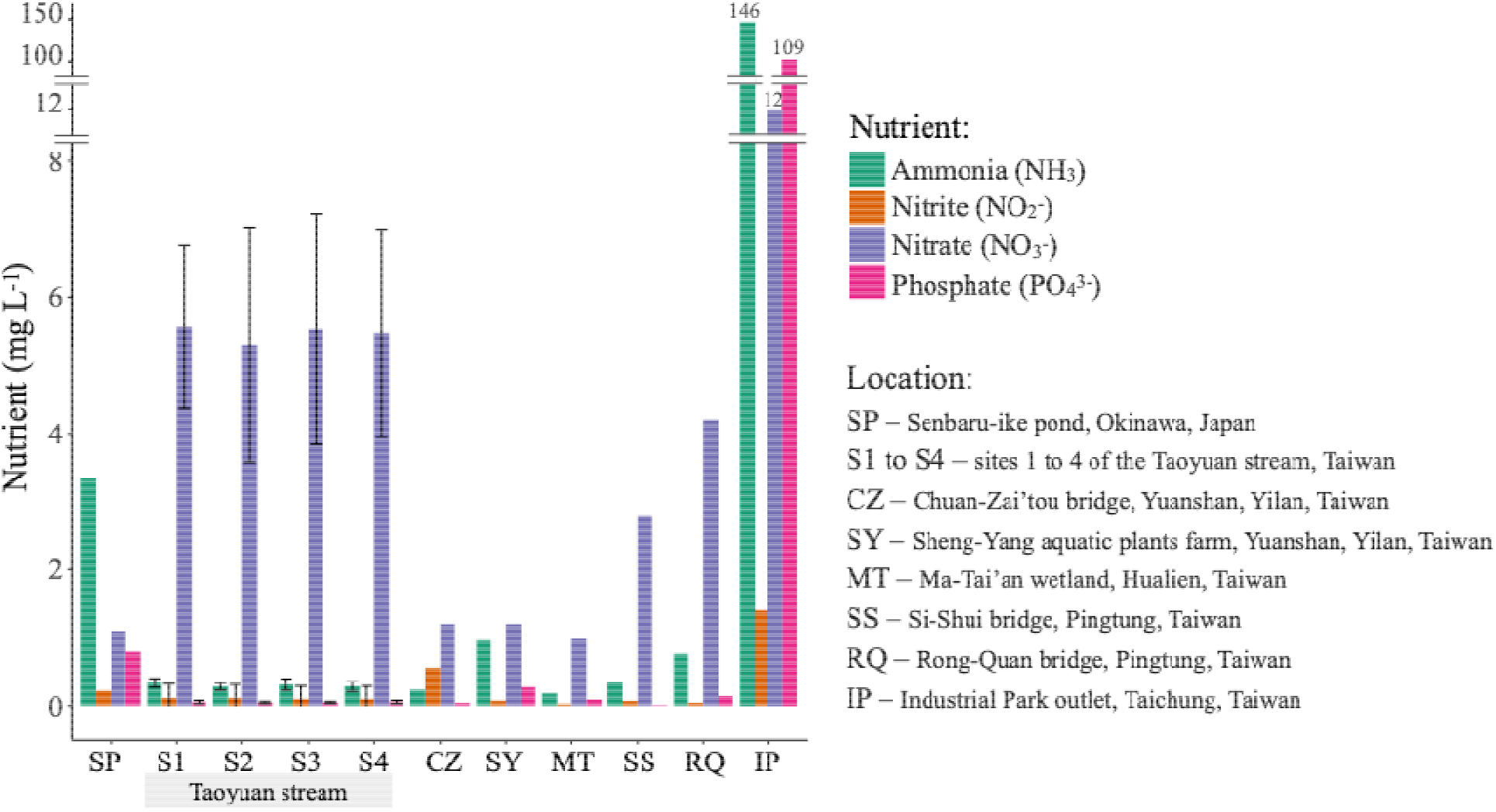
A barplot showing the presence of four nutrients: ammonia, phosphate, nitrite, and nitrate in streams and ponds in Taiwan and Japan where *M. macrospora* was visually detected. Site 1, 2, 3, and 4 (Taoyuan stream) were spots along the Taoyuan stream where monthly surveys were performed. For site 1, 2, 3, and 4, the error bars represent standard deviations. For all the other locations, the level of nutrients was measured once, and therefore error bars could not be computed.

### 3.4 Ecological preferences of *M. macrospora*

Over our 14-month field surveys in the Taoyuan stream, the percent cover of *M. macrospora* at site 3 was significantly higher than the percent cover at site 1 and site 4 (*p* < 0.001; ANOVA and *post hoc* Tukey test; **Figure 6A**; **Tables S4** and **S5**), but marginally significantly higher than the percent cover at site 2 (*p* = 0.0699, ANOVA and *post hoc* Tukey test; **Figure 6A**; **Table S5**). Relative light intensity was significantly higher at site 3, which also had the highest percent cover among the sites, than at the other three sites (*p* < 0.001; one-way ANOVA) (**Figure 6B**; **Table S4** and **S6**). Over the course of our field surveys, the most shaded sites (site 2 and site 4) had a lower light intensity (2 to 2,500 μmol photons m^-2^ s^-1^), and a less shaded site (site 1) had higher light intensity (3 to 8,000 μmol photons m^-2^ s^-1^). In contrast, site 3 was a well-lit, open site that had the highest light intensity (45 to 19,000 μmol photons m^-2^ s^-1^). These results showed that consistent illumination benefits the growth of *M. macrospora*.

**Figure 6.**
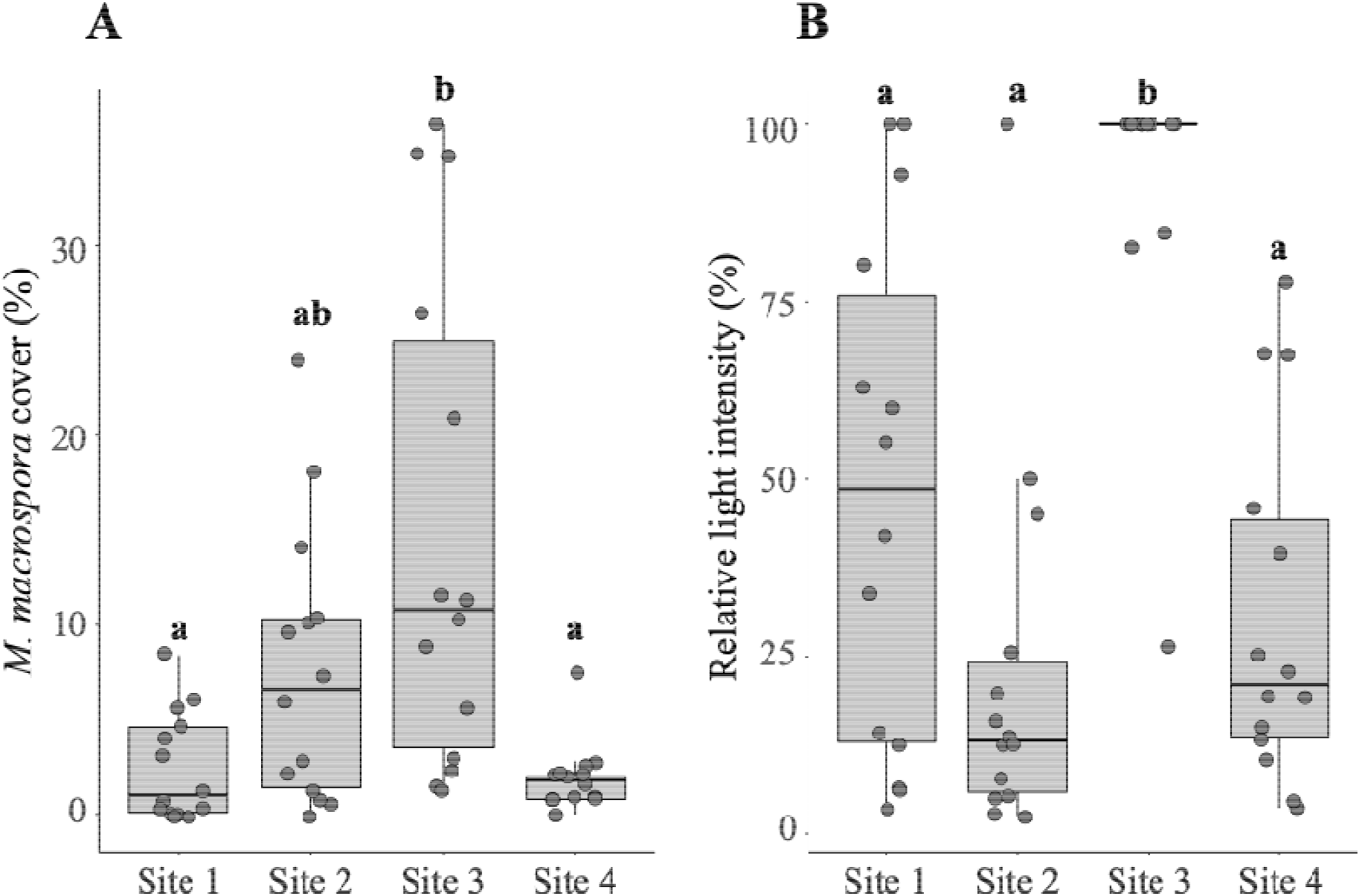
Boxplots showing differences in the percent cover of *M. macrospora* (A) and relative light intensity (B) among four surveyed sites along the Taoyuan stream. Of all the parameters measured, the percent cover of *M. macrospora* and relative light intensity were the only two variables that showed significant variation among the four sites (one-way ANOVA, **Table S4**). *Post hoc* Tukey tests showed significant variation between site 3 and the other three sites, with site 3 having both a higher percent cover of *M. macrospora* and relative light intensity. For each site, single observations are represented as jittered points (*n* = 14).

Next, we performed a stepwise multiple regression analysis to determine which environmental variable was correlated with the percent cover of *M. macrospora*. We conducted an analysis for each of the four sites separately. At sites 1, 2, and 4, majorities of the environmental variables did not significantly explain the temporal change of the percent cover of the algae (**Tables S7** to **S9**); however, ammonia and turbidity were negatively correlated with the algal cover at site 2 and site 3, respectively (**Tables S8** to **S9**). At these three sites, no clear seasonal pattern of algal cover was observed, and the algal cover ranged from nearly 0% to 18% (**Figure S3**). In contrast, at site 3, the percent cover of *M. macrospora* (**Figure 7A**) was positively correlated with water conductivity (a proxy for the concentration of ions) and nitrate (**Figures 7B** and **7D**; **Table S10**), but it was negatively correlated with water temperature, precipitation that accumulated for one week before the survey, ammonia, and turbidity (**Figures 7C** to **7E**; **Table S10**). During the 14-month survey at site 3, the percent cover of *M. macrospora* followed a seasonal pattern (**Figures 7A** and **S3**). It increased up to 36.4% during the winter when the water temperature was lower, and it decreased down to 1.2% during the summer when the water temperature was higher (**Figure 7A**). During our field surveys, the percent cover of other macroalgae was nearly absent (for example, see **Figure 1B**). These observations showed that *M. macrospora* was the most dominant species in the periphyton communities in the surveyed stream.

**Figure 7.**
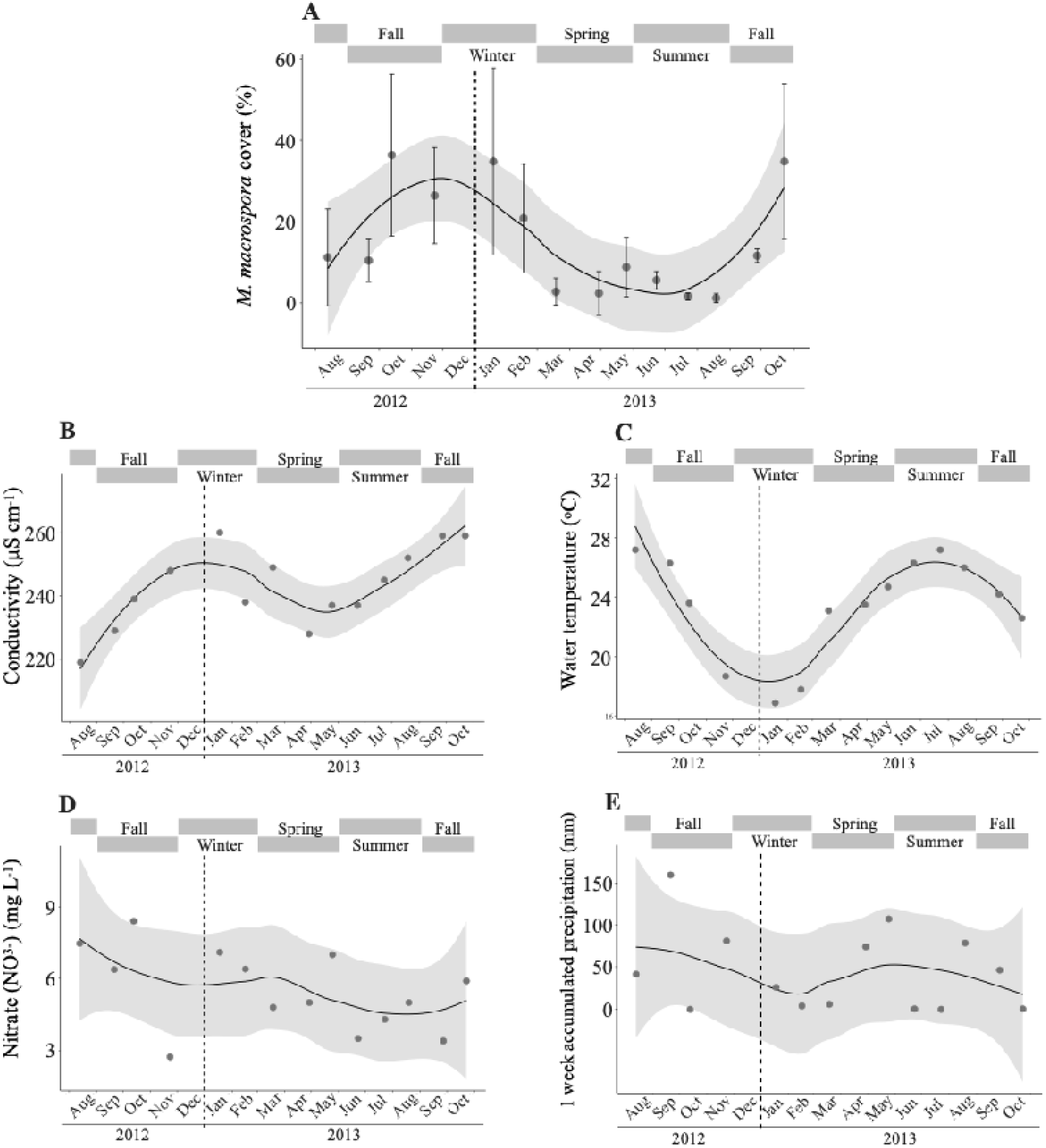
Curve fitting showing the seasonal pattern of percent cover of *M. macrospora* (A) and environmental variables(B-E) at site 3 during a 14-month survey in the Taoyuan stream. Only environmental variables that are mostly significantly correlated with the change of the algal cover are shown based on multiple regression analyses with *p* < 0.01 as follows: conductivity (B), water temperature (C), nitrate level (D), and the amount of precipitation accumulated a week before each survey (E), which was retrieved from Taiwan Weather Bureau (https://www.cwb.gov.tw). The error bars represent standard deviations based on five quadrats along the transect. The shaded area represents a 95% confidence interval around the smoothed regression line. Seasons are indicated as grey bars above each plot.

In addition to the field surveys in the Taoyuan stream, we observed that *M. macrospora* bloomed in the Taichung Industrial Park Outlet. At this site, the percent cover of *M. macrospora* exceeds 50% of all available substrates, such as bamboo shoots, grass shoots, and fishing nets (**Figure S4**). Based on our two time observations spanning nearly eight years (i.e., winter in January, 2014 and summer in September, 2021), this site is warm (25.8-29.7 °C), acidic (pH 5.67-5.83), and highly eutrophic (see above for details).

### 3.5 Observations of other aquarium-associated macrophytes

During the monthly surveys of *M. macrospora* in the Taoyuan stream, we made observations of several macrophytes that are commonly sold in aquaria (**Table 1**, **Figure S5**). Many of the macrophytes were aquatic plants that are commonly transported via the freshwater aquarium trade (*Egeria densa*, *Riccia fluitans, Rotala rotundifolia*, and *Vesicularia mobiana*). Besides *V. mobiana*, these species are not native to Taiwan or Asia, but they were present in the Taoyuan stream, probably due to disposal of aquarium water and unwanted aquatic plants. Also, we occasionally found three freshwater red algae that are commonly present in aquaria: *Compsopogon caeruleus, Nemalionopsis shawii*, and *Sheathia dispersa* (**Table 1**).

**Table 1.**
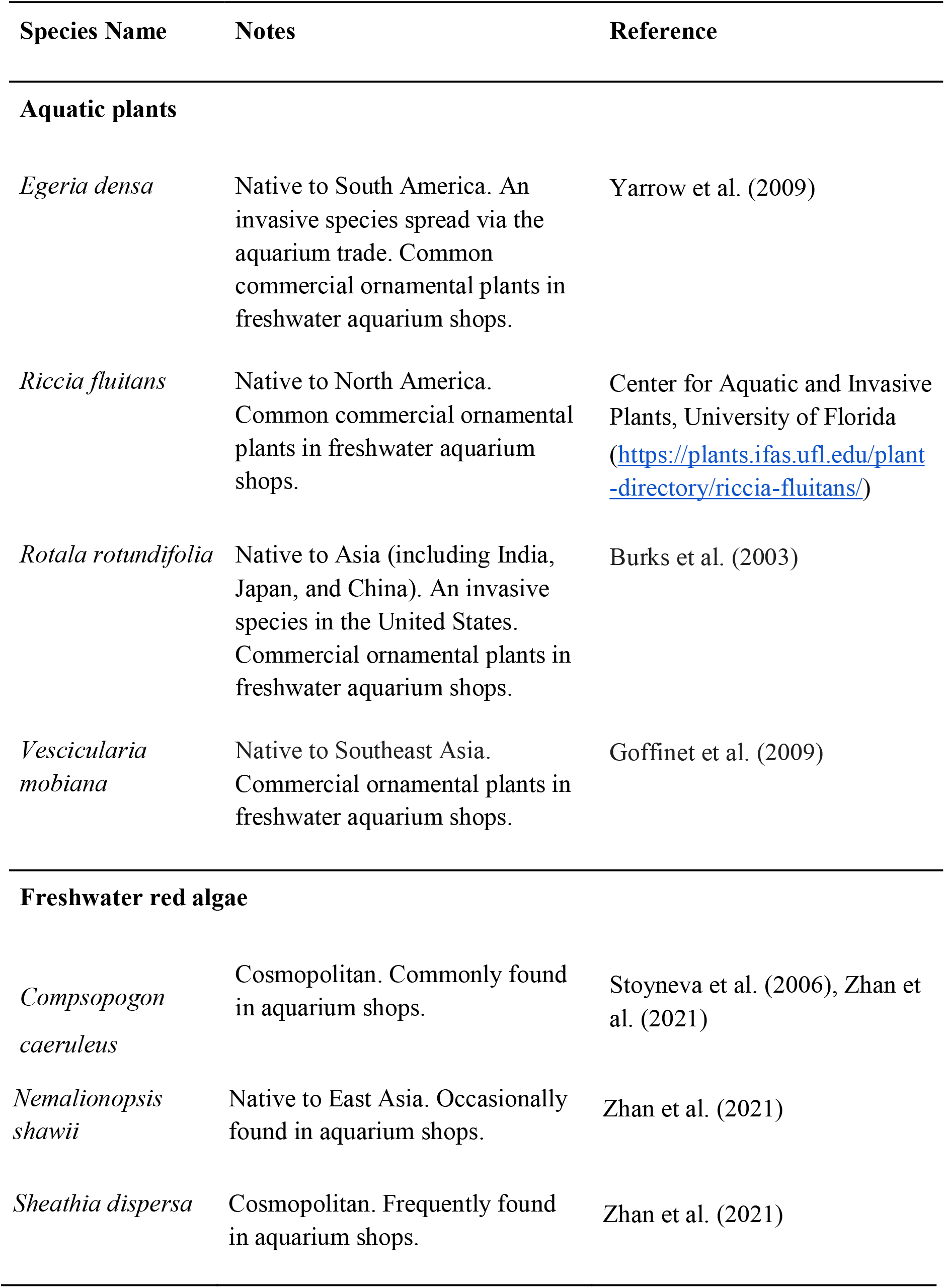
List of aquarium-associated macrophytes co-occurring with *M. macrospora* in the Taoyuan stream. These species are commonly found in aquarium shops. Their *in situ* photos are shown in **Figure S5**.

| Species Name | Notes | Reference |
| --- | --- | --- |
| **Aquatic plants** |  |  |
| *Egeria densa* | Native to South America. An invasive species spread via the aquarium trade. Common commercial ornamental plants in freshwater aquarium shops. | Yarrow et al. (2009) |
| *Riccia fluitans* | Native to North America. Common commercial ornamental plants in freshwater aquarium shops. | Center for Aquatic and Invasive Plants, University of Florida<br>(https://plants.ifas.ufl.edu/plant-directory/riccia-fluitans/) |
| *Rotala rotundifolia* | Native to Asia (including India, Japan, and China). An invasive species in the United States. Commercial ornamental plants in freshwater aquarium shops. | Burks et al. (2003) |
| *Vesicularia mobiana* | Native to Southeast Asia. Commercial ornamental plants in freshwater aquarium shops. | Goffinet et al. (2009) |
| **Freshwater red algae** |  |  |
| *Compsopogon caeruleus* | Cosmopolitan. Commonly found in aquarium shops. | Stoyneva et al. (2006), Zhan et al. (2021) |
| *Nemalionopsis shawii* | Native to East Asia. Occasionally found in aquarium shops. | Zhan et al. (2021) |
| *Sheathia dispersa* | Cosmopolitan. Frequently found in aquarium shops. | Zhan et al. (2021) |

## 4 DISCUSSION

*Montagnia macrospora* is a common freshwater red macroalga in South America (Necchi et al., 2019). It was not until recently that the global aquarium trade was uncovered to be the source of its introduction throughout East Asia (Kato et al., 2009; Zhan et al., 2021). In Taiwan, the earliest record of this alga in the field is dated back to 2005 (Chou et al. 2014). In this study, we conducted a multifaceted ecological assessment of *M. macrospora* in Taiwan via a combination of field surveys, population genetic analysis, and longitudinal environment monitoring.

Our island-wide survey revealed that *M. macrospora* is widespread in the field across Taiwan (about 17% of the sites inspected) in addition to being commonly found in the aquarium shops on the island (Zhan et al. 2021). How could *M. macrospora* have invaded Taiwan and probably other regions of East Asia? One plausible explanation is that the alga is released into the environment via disposal of aquarium water and unwanted aquatic plants by aquarium shop owners and aquarists. In our field observations, *M. macrospora* was often found with other aquarium-associated macrophytes. Another possible explanation is that the alga is dispersed by natural vectors, such as wading waterfowl or rats. Interestingly, viable algae can be found on the feet or in the gut of wading waterfowl and rats (reviewed in Sheath and Hambrook, 1990).

Zhan et al. (2021) have proposed that aquarium shops and aquatic plant farms may serve as *ex situ* reservoirs of *M. macrospora*. How frequently do these local aquarium shops act upon its spread? The data from Zhan et al. (2021) and this study revealed that the sequences of *rbcL* and *cox2-3* are identical across the populations of *M. macrospora* in Taiwan and Okinawa (Japan). This result is consistent with the hypothesis that a founder population of *M. macrospora* was introduced in a single event, probably followed by rapid clonal expansion via asexual reproduction and dispersal among the local aquarium shops or farms. However, we cannot rule out the possibility of repeated introductions of *M. macrospora* from the global trade into local aquarium shops, because the genetic markers used in Zhan et al. (2021) and this study (*rbc*L and *cox2-3*) provide too low resolution to estimate the number of introductions. A microsatellite analysis is needed to test whether there have been multiple, repeated introductions.

Several lines of evidence from this study indicated that *M. macrospora* is an ecological generalist. First, during our surveys, we observed that *M. macrospora* can grow under a broad range of environmental conditions: (1) in cool to warm waters (16 to 30 °C); (2) in acidic to basic waters (pH, 5.67 to 8.34); (3) under shaded (< 65 μmol photons m^-2^ s^-1^) and exposed habitats (> 500 μmol photons m^-2^ s^-1^); (4) in oligotrophic/mesotrophic waters and highly eutrophic waters (e.g., in Senbaru-ike Reservoir and Taichung Industrial Park Outlet); and (5) on abiotic substrates (e.g., plastic tubes, glasses, or cements) and biotic substrates (e.g., submerged shoots of alive or dead grasses, and snail shells). Second, our PCA showed that there was a dramatical niche difference between the native range of *M. macrospora* in South America and the alga’s non-native range in Taiwan, suggesting that *M. macrospora* can cross niche barriers. Third, *M. macrospora* was reported in Taiwan in the Taoyuan stream from 2005 to 2013 and in Taichung Industrial Park Outlet from 2013 to 2021, suggesting that its population can self-sustain and even bloom in new environments for a long period of time. Collectively, these observational data and results show that *M. macrospora* is a generalist and possesses traits observed in other invasive species that are also generalists (Boudouresque and Verlaque, 2002; Evangelista et al., 2008).

Identification of invasive species is important to conservation planning and management, especially in islands such as Taiwan. Islands are key hubs of the global aquarium trade (Padilla and Williams, 2004); thus, they are highly vulnerable to the impacts of invasive species due to a high degree of unique biota (Bellard et al., 2017). What might be the ecological impact of *M. macrospora* on the invaded regions? Having observed that *M. macrospora* can bloom in nonnative habitats, we speculate that one of the ecological impacts of its invasion is outcompeting native macroalgae. Recognizing *M. macrospora* as an invasive species, therefore, paves the way for better understanding its impacts on the invaded regions, such as Taiwan and probably other regions of East Asia.

New species of invasive freshwater macroalgae are rarely reported, unlike species of marine invasive macroalgae that receive a lot more attention (reviewed in Inderjit et al., 2006). The lack of studies focussing on invasive freshwater algae leaves a knowledge gap that may be important to conservation planning and management. The global data analyzed in Zhan et al. (2021) revealed that several species of aquarium-associated freshwater red algae, including *M. macrospora*, had the characteristics of an invasive species (i.e., they exhibit an intercontinental distribution and extremely low genetic diversity), suggesting that there may be more species of invasive freshwater macroalgae introduced via the global aquarium trade that go unrecognized. We encourage researchers who specialize in conservation biology and management with an interest in algae ecology to collaborate with phycologists in the future to improve our understanding of the role that invasive freshwater algae play on introduced habitats.

## ACKNOWLEDGEMENTS

We thank Tsai-Yin Hsieh and Ganies Riza Aristya (Tunghai University) for their help with the nutrient measurements and photography. We also thank Dr. Shoichiro Suda (University of Ryukyus) for helping the collection of the water sample in the Senbaru-ike pond in Okinawa. SF was financially supported by the MOST postdoctoral fellowships of the Ministry of Science and Technology (MOST), Taiwan (MOST109-2811-B-029-501 and MOST110-2811-B-029-002). This study was supported by grants from MOST, Taiwan to SLL (MOST108-2624-B-029-005-MY3).

## AUTHOR CONTRIBUTIONS

**Silvia Fontana:** Conceptualization (Equal); Data curation (Equal); Formal analysis (Equal); Methodology (Equal); Software (Equal); Visualization (Equal); Writing-original draft (Equal); Writing-review & editing (Equal). **Lan-Wei Yeh:** Data curation (Equal); Formal analysis (Equal); Methodology (Equal); Software (Equal); Visualization (Equal); Writing-original draft (Equal); Writing-review & editing (Equal).. **Shing Hei Zhan:** Conceptualization (Equal); Validation (Equal); Visualization (Equal); Writing-review & editing (Equal). **Shao-Lun Liu:** Conceptualization (Equal); Data curation (Equal); Formal analysis (Equal); Funding acquisition (Equal); Investigation (Equal); Methodology (Equal); Supervision (Equal); Validation (Equal); Visualization (Equal); Writing-review & editing (Equal).

## DATA AVAILABILITY STATEMENT

The *cox2-3* sequence newly generated in this study is available at NCBI GenBank under accession numbers: OK442460.

