## Supplemental Information for "A multifaceted ecological assessment reveals the invasion of the freshwater red macroalga *Montagnia macrospora* (Batrachospermales, Rhodophyta) in Taiwan"

**Table S1.** Bioclimatic variables and elevation data of the sampling sites in Taiwan and South America where *Montagnia macrospora* was found**.** Occurrence data are obtained from Vis et al. (2008), Zhan et al. (2021), and this study.

| **Site** | **Longitude** | **Latitude** | **Elevation** | **Bio02** | **Bio03** | **Bio08** | **Bio09** | **Bio13** | **Bio14** | **Bio15** | **Bio18** | **Bio19** |
| --- | --- | --- | --- | --- | --- | --- | --- | --- | --- | --- | --- | --- |
| **Taiwan (field only)** | | | | | | | | | | | | |
| 1 | 121.2315 | 24.88541 | 209 | 6.3 | 3.2 | 26.5 | 16.4 | 290 | 66 | 40 | 791 | 349 |
| 2 | 121.7053 | 24.72818 | 21 | 6.1 | 3.3 | 23.3 | 18.3 | 429 | 121 | 39 | 844 | 462 |
| 3 | 121.4088 | 23.65814 | 210 | 7 | 4.1 | 26.6 | 17.9 | 353 | 62 | 54 | 762 | 214 |
| 4 | 120.5788 | 22.58431 | 20 | 7.8 | 4.7 | 27.6 | 20.9 | 719 | 13 | 105 | 1784 | 58 |
| 5 | 120.5135 | 22.71915 | 33 | 8.1 | 4.7 | 27.6 | 20.7 | 611 | 14 | 106 | 1591 | 57 |
| 6 | 120.6014 | 24.10534 | 23 | 7.4 | 3.7 | 27.6 | 20.9 | 315 | 12 | 83 | 898 | 113 |
| 7 | 120.5619 | 24.21435 | 39 | 7.3 | 3.7 | 27.5 | 20.7 | 307 | 10 | 83 | 860 | 117 |
| 8 | 121.6983 | 24.72999 | 38 | 6.1 | 3.3 | 25.7 | 18.3 | 432 | 122 | 39 | 866 | 463 |
| **South America** | | | | | | | | | | | | |
| 1 | -49.33333 | -22.81667 | 602 | 12.8 | 6.5 | 23.1 | 17.2 | 200 | 29 | 55 | 569 | 146 |
| 2 | -44.94254 | -23.35298 | 33 | 9.4 | 5.6 | 25.5 | 20.1 | 315 | 92 | 41 | 929 | 297 |
| 3 | -44.20889 | -21.30028 | 1036 | 11.9 | 6.5 | 21.5 | 16.5 | 304 | 15 | 78 | 665 | 72 |
| 4 | -52.64583 | 4.585556 | 55 | 8.2 | 7.6 | 25.5 | 26.1 | 532 | 90 | 46 | 360 | 1144 |
| 5 | -53.58389 | 5.623611 | 0 | 8.1 | 7.8 | 26.6 | 27.2 | 363 | 49 | 51 | 201 | 561 |
| 6 | -41.05 | -20.31667 | 966 | 11.1 | 6.4 | 20.5 | 16.9 | 216 | 33 | 56 | 453 | 140 |
| 7 | -53.06417 | 5.3025 | 64 | 7.9 | 7.7 | 25.8 | 26.3 | 453 | 54 | 50 | 245 | 789 |
| 8 | -67.75361 | -16.25306 | 2268 | 13.4 | 7.5 | 18.1 | 15.7 | 201 | 15 | 68 | 445 | 64 |
| 9 | -47.78333 | -22.15 | 782 | 11 | 6.3 | 21.8 | 16.7 | 255 | 24 | 70 | 622 | 109 |
| 10 | -60.04889 | -2.709722 | 50 | 8.4 | 8.2 | 27 | 27.6 | 281 | 86 | 34 | 395 | 723 |
| 11 | -59.70361 | -2.699722 | 34 | 8.5 | 8.2 | 27 | 27.8 | 287 | 99 | 35 | 370 | 777 |
| 12 | -52.42556 | 4.836944 | 15 | 7.7 | 7.5 | 26.1 | 27.1 | 542 | 61 | 51 | 273 | 1046 |
| 13 | -53.04755 | 5.062967 | 30 | 8 | 7.8 | 25.9 | 26.4 | 446 | 70 | 47 | 280 | 841 |
| 14 | -52.40667 | 4.794444 | 14 | 7.7 | 7.4 | 26.1 | 27.1 | 555 | 67 | 50 | 298 | 1091 |
| 15 | -52.72843 | 5.149805 | 11 | 7.8 | 7.7 | 26 | 26.4 | 471 | 49 | 53 | 252 | 877 |
| 16 | -53.97463 | 5.488652 | 21 | 8.3 | 7.8 | 26.5 | 27.3 | 353 | 86 | 38 | 333 | 599 |

Bio02, mean of monthly temperature (max temp − min temp), or the mean diurnal range; Bio03, isothermality ((mean diurnal range/temperature annual range)*100); Bio08, mean temperature of wettest quarter; Bio09, mean temperature of driest quarter; Bio13, precipitation of wettest month; Bio14, precipitation of driest month; Bio15, precipitation seasonality (coefficient of variation); Bio18, precipitation of warmest quarter; Bio19, precipitation of coldest quarter.

**Table S2.** Collection locations and dates of the samples of *Montagnia macrospora* used to sequence the *cox2-3* marker. Samples collected from this study (Taiwan and Japan) were integrated with sequences retrieved from GenBank (South America).

| GPS (Lat/lon) | Location (Voucher Number) | Habitat | Collection date | Haplotype/Reference | GenBank Accession |
| --- | --- | --- | --- | --- | --- |
| 24.870909/121.22406 | "ChangJiang" Aquarium Shop, Zhongxing Road, Longtan Township, Taoyuan, Taiwan (THU.007) | aquarium | 29/04/2013 | -/this study | OK442460 |
| 24.870909/121.22406 | "ChangJiang" Aquarium Shop, Zhongxing Road, Longtan Township, Taoyuan, Taiwan (THU.008) | aquarium | 29/04/2013 | -/this study | The same as THU.007. |
| 24.177590/120.598283 | Home aquarium tank, Taichung, Taiwan (THU.010) | aquarium | 23/06/2013 | -/this study | The same as THU.007. |
| 23.956610/120.567329 | Yulaike Aquarium Shop, Yuanlin, Changhua, Taiwan (THU.014) | aquarium | 26/06/2013 | -/this study | The same as THU.007. |
| 23.956610/120.567329 | Yulaike Aquarium Shop, Yuanlin, Changhua, Taiwan (THU.015) | aquarium | 26/06/2013 | -/this study | The same as THU.007. |
| 22.642742/120.329090 | Gaojianduan Aquarium Shop, Kaohsiung, Taiwan (THU.039) | aquarium | 26/06/2013 | -/this study | The same as THU.007. |
| 22.672366/120.297929 | Yung Shin Aquarium Shop, Kaohsiung, Taiwan (THU.043) | aquarium | 26/06/2013 | -/this study | The same as THU.007. |
| 26.694158/127.877932 | Aquarium tank at Makeman Newman, Nishihara-ch, Okinawa, Japan (THU.078) | aquarium | 31/08/2013 | -/this study | The same as THU.007. |
| 24.885412/121.231454 | Spring-fed Irrigation ditch in front of "HuangNiTangFuDe" Temple, Longping Road, Longtan, Taoyuan, Taiwan (THU.125) | spring | 13/08/2012 | -/this study | The same as THU.007. |
| 24.885412/121.231454 | Spring-fed Irrigation ditch in front of "HuangNiTangFuDe" Temple, Longping Road, Longtan, Taoyuan, Taiwan (THU.126) | spring | 13/08/2012 | -/this study | The same as THU.007. |
| 24.885412/121.231454 | Spring-fed Irrigation ditch in front of "HuangNiTangFuDe" Temple, Longping Road, Longtan, Taoyuan, Taiwan (THU.127) | spring | 13/08/2012 | -/this study | The same as THU.007. |
| 24.885412/121.231454 | Spring-fed Irrigation ditch in front of "HuangNiTangFuDe" Temple, Longping Road, Longtan, Taoyuan, Taiwan (THU.128) | spring | 13/08/2012 | -/this study | The same as THU.007. |
| 24.885412/121.231454 | Spring-fed Irrigation ditch in front of "HuangNiTangFuDe" Temple, Longping Road, Longtan, Taoyuan, Taiwan (THU.129) | spring | 13/08/2012 | -/this study | The same as THU.007. |
| 24.885412/121.231454 | Spring-fed Irrigation ditch in front of "HuangNiTangFuDe" Temple, Longping Road, Longtan, Taoyuan, Taiwan (THU.130) | spring | 13/08/2012 | -/this study | The same as THU.007. |
| 24.885412/121.231454 | Spring-fed Irrigation ditch in front of "HuangNiTangFuDe" Temple, Longping Road, Longtan, Taoyuan, Taiwan (THU.147) | spring | 13/10/2012 | -/this study | The same as THU.007. |
| 24.885412/121.231454 | Spring-fed Irrigation ditch in front of "HuangNiTangFuDe" Temple, Longping Road, Longtan, Taoyuan, Taiwan (THU.148) | spring | 13/10/2012 | -/this study | The same as THU.007. |
| 24.885412/121.231454 | Spring-fed Irrigation ditch in front of "HuangNiTangFuDe" Temple, Longping Road, Longtan, Taoyuan, Taiwan (THU.149) | spring | 13/10/2012 | -/this study | The same as THU.007. |
| 24.885412/121.231454 | Spring-fed Irrigation ditch in front of "HuangNiTangFuDe" Temple, Longping Road, Longtan, Taoyuan, Taiwan (THU.150) | spring | 13/10/2012 | -/this study | The same as THU.007. |
| 24.885412/121.231454 | Spring-fed Irrigation ditch in front of "HuangNiTangFuDe" Temple, Longping Road, Longtan, Taoyuan, Taiwan (THU.151) | spring | 13/10/2012 | -/this study | The same as THU.007. |
| 22.719145/120.513516 | Yuquan Village, Pingtung, Taiwan (THU.264) | stream | 25/01/2014 | -/this study | The same as THU.007. |
| 22.719145/120.513516 | Yuquan Village, Pingtung, Taiwan (THU.319) | stream | 25/01/2014 | -/this study | The same as THU.007. |
| 22.719145/120.513516 | Yuquan Village, Pingtung, Taiwan (THU.321) | stream | 25/01/2014 | -/this study | The same as THU.007. |
| 26.247582/127.765142 | An artificial reservoir pond, Senbaru-ike, at University of the Ryukyus, Okinawa, Japan (THU.73) | pond | 10/09/2013 | -/this study | The same as THU.007. |
| 24.105335/120.601382 | Taichung Industrial Park Outlet, Taichung, Taiwan (THU.476) | stream | 27/12/2013 | -/this study | The same as THU.007. |
| 5.772777778/-53.51638889 | L'Organabo Riviere on road N1 at public access, west of Iracoubo, French Guiana | stream | 19/08/2002 | I/Vis et al. (2008) | EU106072 |
| -23.35/-44.94472222 | Rio Puruba, Sao Paulo State, Brazil | stream | 15/12/2001 | J/Vis et al. (2008) | EU106073 |
| -2.709722222/-60.04888889 | Taruma on BR 174 km 31, near Manaus, Brazil | stream | 20/08/2004 | K/Vis et al. (2008) | EU106074 |
| -2.699722222/-59.70361111 | Rio Preto da Eva, near Manaus, Brazil | stream | 20/08/2004 | L/Vis et al. (2008) | EU106075 |
| -2.709722222/-60.04888889 | Taruma on BR 174 km 31, near Manaus, Brazil | stream | 20/08/2004 | M/Vis et al. (2008) | EU106076 |
| -2.699722222/-59.70361111 | Rio Preto da Eva, near Manaus, Brazil | stream | 20/08/2004 | N/Vis et al. (2008) | EU106077 |
| -2.699722222/-59.70361111 | Rio Preto da Eva, near Manaus, Brazil | stream | 20/08/2004 | O/Vis et al. (2008) | EU106078 |
| -2.709722222/-60.04888889 | Taruma on BR 174 km 31, near Manaus, Brazil | stream | 20/08/2004 | P/Vis et al. (2008) | EU106079 |

**Table S3.**  Eigenvectors of the first three principal components (PCs) of the principal component analysis that involve the ten climatic variables in **Table S1**.

| Bioclimatic variables | Factor Loading | |
| --- | --- | --- |
|  | **PC1** | **PC2** |
| Elevation | **0.384** | -0.284 |
| Bio02 | **0.337** | **-0.33** |
| Bio03 | -0.147 | **-0.424** |
| Bio08 | **-0.328** | **0.34** |
| Bio09 | **-0.409** | -0.101 |
| Bio13 | -0.223 | **0.321** |
| Bio14 | **-0.329** | -0.106 |
| Bio15 | 0.273 | **0.336** |
| Bio18 | 0.183 | **0.487** |
| Bio19 | **-0.42** | -0.2 |

Bio02, mean of monthly temperature (max temp − min temp), or the mean diurnal range; Bio03, isothermality ((mean diurnal range/temperature annual range)*100); Bio08, mean temperature of wettest quarter; Bio09, mean temperature of driest quarter; Bio13, precipitation of wettest month; Bio14, precipitation of driest month; Bio15, precipitation seasonality (coefficient of variation); Bio18, precipitation of warmest quarter; Bio19, precipitation of coldest quarter.

**Table S4.** Results of independent one-way ANOVA testing variations in the percent cover of *Montagnia macrospora* and collected environmental parameters among the four sites surveyed during the study (**p* < 0.05, ***p* < 0.01, ****p* <0.001).

| Variable | Df | Sum Sq | Mean Sq | F value | *p* value |
| --- | --- | --- | --- | --- | --- |
| Algae coverage (%) | 3 | 1,529 | 509.69 | 8.57 | 0.0001013*** |
| Relative light intensity (%) | 3 | 40,677 | 13,559 | 17.93 | 4.059e-08*** |
| pH | 3 | 0.48 | 0.16 | 0.73 | 0.54 |
| Temperature | 3 | 1.04 | 0.35 | 0.03 | 0.99 |
| Conductivity | 3 | 110.1 | 36.69 | 0.29 | 0.84 |
| Turbidity | 3 | 39.17 | 13.06 | 0.37 | 0.78 |
| Ammonia (NH_3_) | 3 | 0.03 | 0.01 | 1.96 | 0.13 |
| Phosphate (PO_4_^3-^) | 3 | 0 | 0 | 0.51 | 0.67 |
| Nitrite (NO_2_^-^) | 3 | 0 | 0 | 0.03 | 0.99 |
| Nitrate (NO_3_^-^) | 3 | 0.55 | 0.18 | 0.08 | 0.97 |

**Table S5.** Results of a multi-way comparison of the percent cover of *Montagnia macrospora* among the four sites surveyed during the study using *post hoc* Tukey test (**p* < 0.05, ***p* < 0.01, ****p* <0.001).

| Linear hypotheses | Estimate | Std. Error | t value | *p* value |
| --- | --- | --- | --- | --- |
| site 2 – site 1 = 0 | 5.114 | 2.915 | 1.754 | 0.3068 |
| site 3 – site 1 = 0 | 12.429 | 2.915 | 4.264 | <0.001 *** |
| site 4 – site 1 = 0 | -0.600 | 2.915 | -0.206 | 0.9969 |
| site 3 – site 2 = 0 | 7.314 | 2.915 | 2.509 | 0.0699 |
| site 4 – site 2 = 0 | -5.714 | 2.915 | -1.960 | 0.2163 |
| site 4 – site 3 = 0 | -13.029 | 2.915 | -4.469 | <0.001 *** |

**Table S6.** Results of a multi-way comparison of the relative light intensity (%) among the four sites surveyed during the study using *post hoc* Tukey test (**p* < 0.05, ***p* < 0.01, ****p* < 0.001).

| Linear hypotheses | Estimate | Std. Error | t value | *p* value |
| --- | --- | --- | --- | --- |
| site 2 – site 1 = 0 | -25.122 | 10.395 | -2.417 | 0.0865 |
| site 3 – site 1= 0 | 44.571 | 10.395 | 4.288 | <0.001 *** |
| site 4 – site 1= 0 | -17.010 | 10.395 | -1.636 | 0.3676 |
| site 3 – site 2= 0 | 69.692 | 10.395 | 6.705 | <0.001 *** |
| site 4 – site 2= 0 | 8.112 | 10.395 | 0.780 | 0.8630 |
| site 4 – site 3= 0 | -61.581 | 10.395 | -5.924 | <0.001 *** |

**Table S7.** Results of a multiple regression analysis followed by stepwise selection, applied to the survey on site 1. Shown are the variables (out of the environmental variables: water temperature, pH, conductivity, turbidity, and accumulated precipitation one week before the survey, and four nutrients: NH_3_, PO_4_^3-^, NO_2_^-^, and NO_3_^-^) selected to fit the best-performing model to explain algal percent cover. F-statistic *p* value = 0.1706.

| Site 1 | Estimate | Std. Error | t value | *p* value |
| --- | --- | --- | --- | --- |
| (Intercept) | 1.62 | 0.38 | 0.24 | 0.82 |
| Phosphate (PO_4_^3-^) | 42.06 | 28.85 | 1.46 | 0.17 |

**Table S8.** Results of a multiple regression analysis followed by stepwise selection, applied to the survey on site 2. Shown are the variables (out of the environmental variables: water temperature, pH, conductivity, turbidity, and accumulated precipitation one week before the survey, and four nutrients: NH_3_, PO_4_^3-^, NO_2_^-^, and NO_3_^-^) selected to fit the best-performing model to explain algal percent cover. F-statistic *p* value = 0.06181.

| Site 2 | Estimate | Std. Error | t value | *p* value |
| --- | --- | --- | --- | --- |
| (Intercept) | -40.16 | 31.32 | -1.28 | 0.230 |
| pH | 10.45 | 4.59 | 2.28 | 0.050 |
| Ammonia  (NH_3_) | -83.53 | 34.84 | -2.4 | 0.040* |
| Nitrite (NO_2_^-^) | -18.9 | 14.69 | -1.29 | 0.230 |

**Table S9.** Results of a multiple regression analysis followed by stepwise selection, applied to the survey on site 4. Shown are the variables (out of the environmental variables: water temperature, pH, conductivity, turbidity, and accumulated precipitation one week before the survey, and four nutrients: NH_3_, PO_4_^3-^, NO_2_^-^, and NO_3_^-^) selected to fit the best-performing model to explain algal percent cover. F-statistic *p* value = 0.004828.

| Site 4 | Estimate | Std. Error | t value | *p* value |
| --- | --- | --- | --- | --- |
| (Intercept) | 10.2 | 9.42 | 1.08 | 0.300 |
| pH | 1.7 | 0.88 | 1.93 | 0.080 |
| Turbidity | -0.18 | 0.06 | -2.93 | 0.020* |
| Nitrate (NO_3_^-^) | 0.47 | 0.23 | 2.08 | 0.060︎ |

**Table S10.** Results of a multiple regression analysis followed by stepwise selection, applied to the survey on site 3. Shown are the variables (out of the environmental variabales: water temperature, pH, conductivity, turbidity, and accumulated precipitation one week before the survey, and four nutrients: NH_3_, PO_4_^3-^, NO_2_^-^, and NO_3_^-^) selected to fit the best-performing model to explain algal percent cover. F-statistic *p* value = 0.002384.

| Site 3 | Estimate | Std. Error | t value | *p* value |
| --- | --- | --- | --- | --- |
| (Intercept) | 94.4 | 84.28 | 1.12 | 0.330 |
| Water temperature | -2.37 | 0.38 | -6.17 | 0.004** |
| pH | -8.09 | 5.75 | -1.41 | 0.230 |
| Conductivity | 1.48 | 0.21 | 6.98 | 0.002** |
| Turbidity | -2.44 | 0.63 | -3.85 | 0.020* |
| Ammonia (NH_3_) | -92.8 | 26.55 | -3.5 | 0.020* |
| Phosphate (PO_4_^3-)^ | -186.3 | 87.84 | -2.12 | 0.100 |
| Nitrite (NO_2_^-^) | -24.89 | 9.67 | -2.57 | 0.060︎ |
| Nitrate (NO_3_^-^) | 3.85 | 0.67 | 5.77 | 0.004** |
| Accumulated precipitation 1 week before collection | -0.23 | 0.03 | -7.21 | 0.002** |

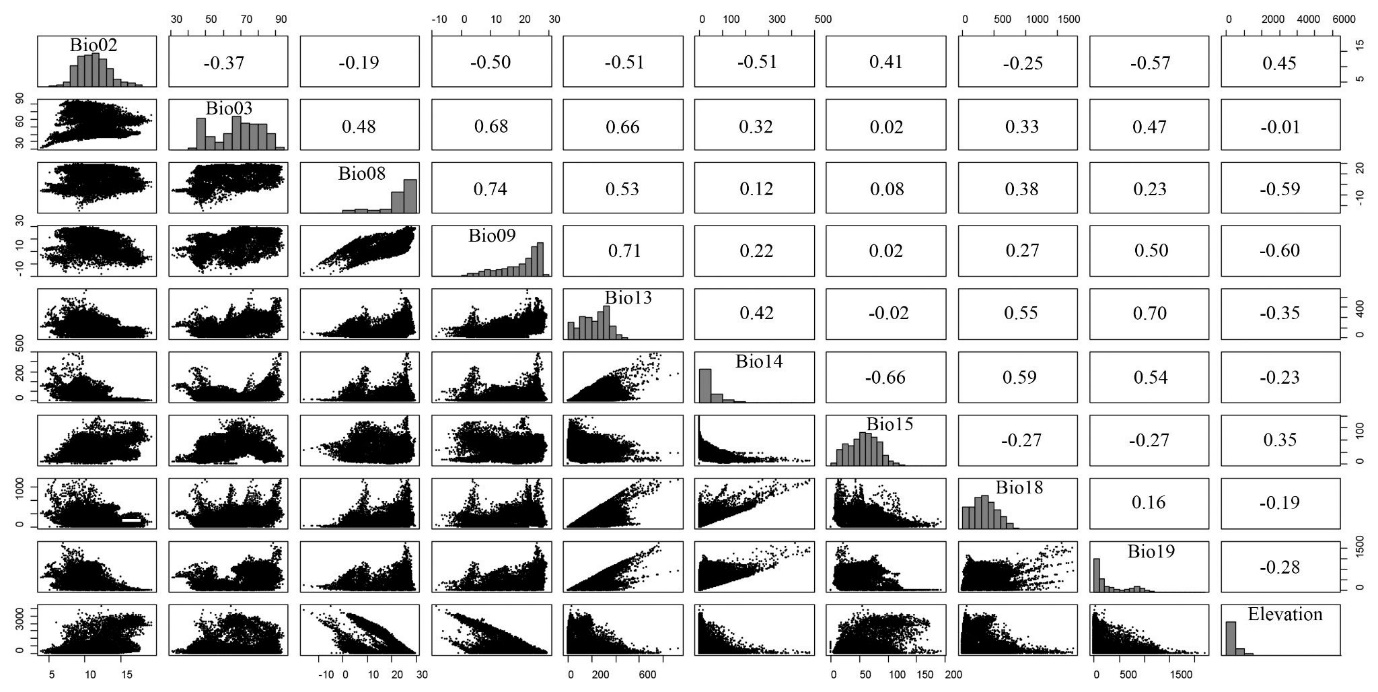

**Figure S1.** Pairwise correlation analysis of the ten climatic variables selected after a multicollinearity test.

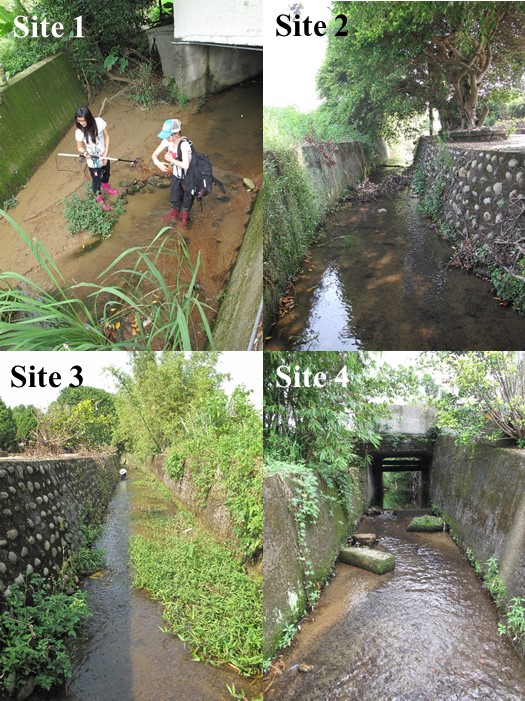

**Figure S2.** Examples showing the four sites along the Taoyuan stream during the 14-month field survey. Site 3 enjoyed the most sunlight, and the other three sites showed different degrees of tree canopy shading. Site 1 was muddier than the other sites, sites 2 and 3 had cemented stream beds, and site 4 had silts in its bed.

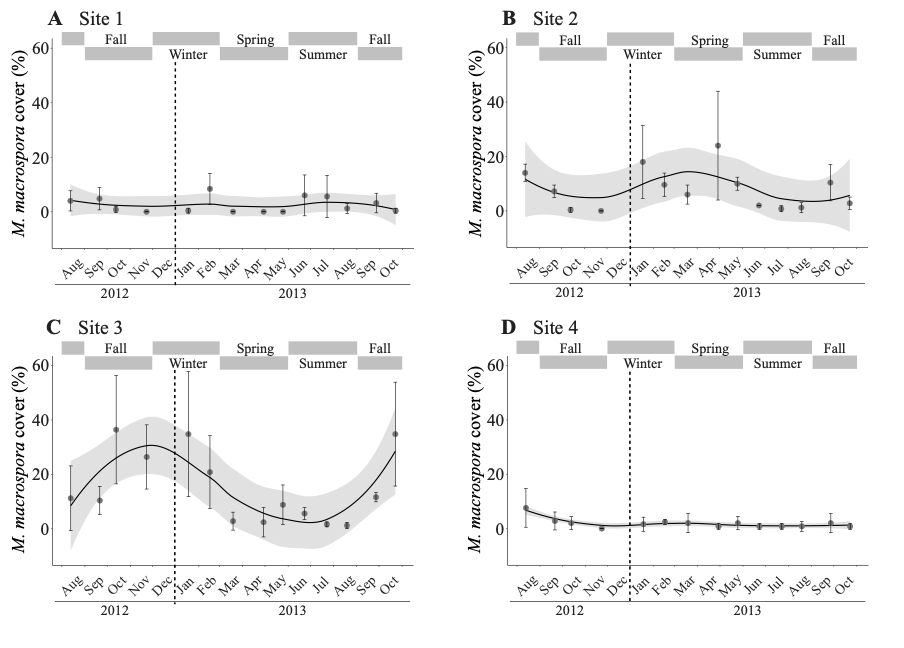

**Figure S3.** Percentage coverage of *M. macrospora* in the four sites during the monthly survey in a stream near Taoyuan (Taiwan): site 1 (A), site 2 (B), site 3 (C), and site 4 (D). The data are the average of five quadrats along each transect (site). The error bars represent standard deviations. The 95% confidence intervals are shaded around the smoothed regression lines. Meteorological seasons are highlighted in each panel.

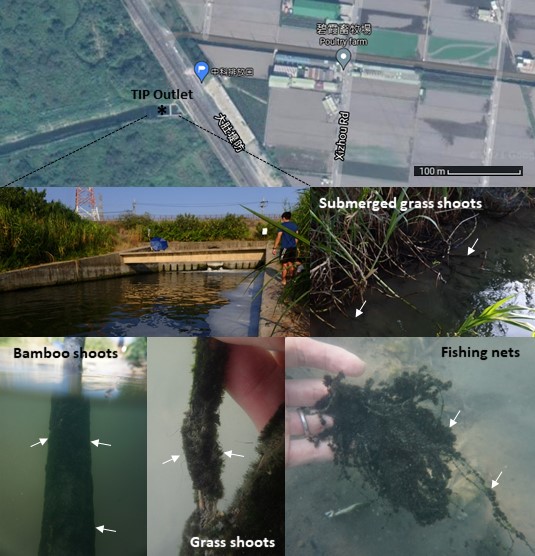

**Figure S4.** Photos showing the overgrowth of *Montagnia macrospora* in all available substrates in the Taichung Industrial Park Outlet, Taiwan (24°08'01.2"N 120°32'23.3"E) in September, 2021.

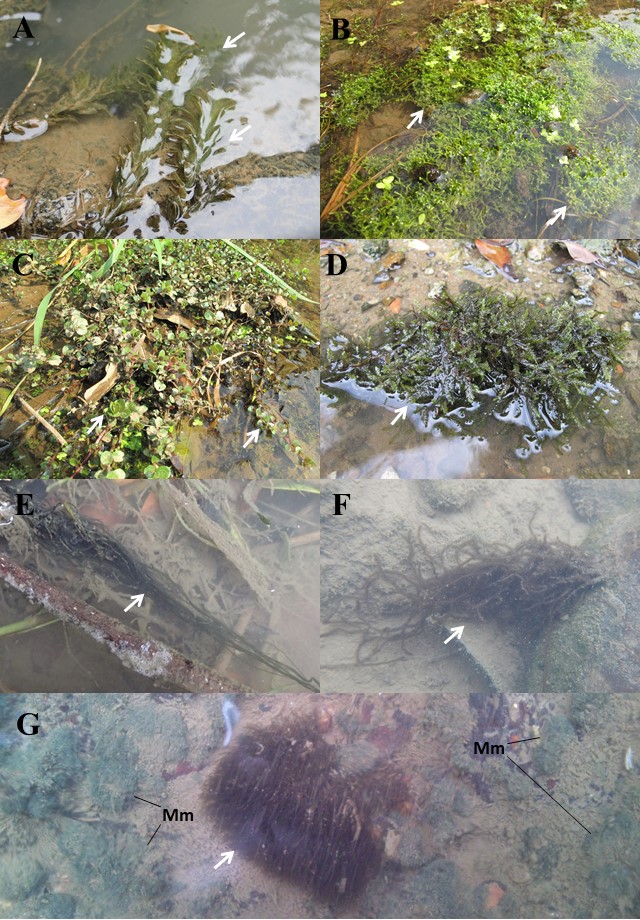

**Figure S5.** *In situ* photos showing aquatic plants (white arrows in A-D) and freshwater red algae (white arrows in E-G) that are commonly found in the aquarium trade. (A) *Egeria densa*. (B) *Riccia fluitans*. (C) *Rotala rotundifolia*. (D) *Vescicularia mobiana* (also known as Java moss). (E) *Compsopogon caeruleus*. (F) *Nemalionopsis shawii*. (G) *Sheathia dispersa* intermixing with the most dominant benthic macroalga, *Montagnia macrospora* (Mm). Detailed information about the occurrence of each species in the aquarium shops are listed in **Table 1**.
